# Impact of life origins on metabolism

**DOI:** 10.1101/2024.06.07.597902

**Authors:** Alexei Vazquez

## Abstract

Living organisms are defined by self-replication and self-confinement. We expect these two properties to shape the metabolic capabilities of cells. Here I demonstrate that the maximum growth rate of cells is, in a first approximation, the geometric mean between the maximum rate of ribosome self-replication and the maximum rate of macromolecular synthesis allowed by the interior volume defined by the cell membrane. I also show how these constraints are buried into the biomass compositions of flux balance models.

## 1. Introduction

Self-replication and self-confinement are at the root of the origins of life [1]. Self-replication allows cells to make copies of themselves. Self-confinement enforces the separation between self and the environment. The action of natural selection under these two key properties should be reflected in the metabolic efficiency of cells.

Studies aiming to identify the minimal components the last universal common ancestor (LUCA) concord on the presence of ribosomes and a chemiostatic membrane [2, 3, 4]. The ribosomes are the cell machinery devoted to protein synthesis and they are composed of proteins and RNA. Because the ribosomes synthesize their own proteins, the ribosomes engage in cell replication. Not surprisingly, the ribosomes are optimized for self-replication [5, 6, 7].

The presence of a cell membrane since LUCA has been studied from the point of view of its chemiostatic nature and the membrane bound ATP synthase [3]. What is less appreciated is that the confinement associated with the cell membrane imposes a molecular crowding constraint on cells metabolism. In growing cells the cell membrane area scales in proportion to the cell volume [8, 9]. The link between this observation, molecular crowding and the limits of cell growth are investigated here for the first time.

## 2. Self-replication limit

The consequences of self-replication on metabolic efficiency has been extensively studied [10, 11, 5, 6]. A key example is the self-replicative nature of protein synthesis, whereby the protein synthesis machinery ℰ_*P*_, the ribosomes, is itself made or proteins. Suppose we drop *E*_*P*_ (0) ribosomes in a broth containing ribosomal RNA, amino acids, ATP and everything else needed for protein synthesis. The ribosomes will undergo the synthesis of more ribosomes. If we denote by *E*_*P*_ (*t*) the number of ribosomes at a given time *t*, then the evolution in time of the number of ribosomes will follow the first order differential equation

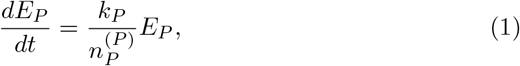

where *k*_*P*_ is the rate of protein synthesis in units of amino acids added to polypeptide chains per unit of ribosome and 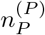 is the number of protein precursors (amino acids) per protein synthesis machinery (ribosome). The solution to this differential equation is the exponential growth 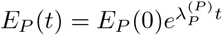, with a replication rate

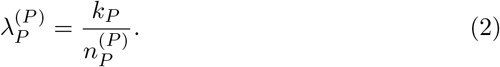

I will call the latter the protein synthesis self-replication limit.

## 3. Confinement limit

The impact of confinement on metabolic efficiency is less appreciated. I have pushed forward the idea that molecular crowding, a natural outcome of confinement, limits cell metabolism [12, 13, 14, 15, 16]. I am not convinced that we fully understand its *modus operandi*. Here I present a more intuitive approach. The key feature of confinement is the existence of a cell membrane separating self from the environment. From the point of view of metabolism, this membrane is synthesized by some protein machinery ℰ_*M*_, the proteomic set containing all proteins associated with the cytoplasmic membrane synthesis. If we denote by *E*_*M*_ the number of ℰ_*M*_ units in the cell, then the cell membrane area should grow according to the differential equation

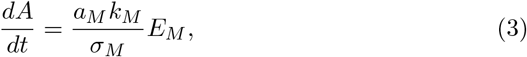

where *k*_*M*_ is the rate of membrane precursor synthesis per unit of ℰ_*M*_, *a*_*M*_ is the membrane precursor molar area and *σ*_*M*_ is the area fraction occupied by the membrane precursor. Assuming that the cell and the ℰ_*M*_ content are growing exponentially with a rate *λ*, then the solution to this equation is

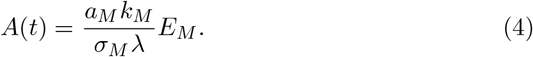

Now the crowding constraint kicks in. If the cell volume scale as *V* = *lA*, where *l* is the volume to surface ratio, then the cell volume fraction occupied by ℰ_ℳ_ is given by

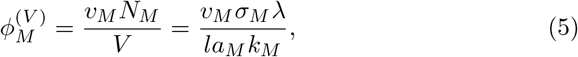

where *v*_*M*_ is the molar volume of ℰ_ℳ_. This volume fraction cannot exceed the maximum packing density Φ_max_ and, therefore, equation (5) implies the maximum growth rate

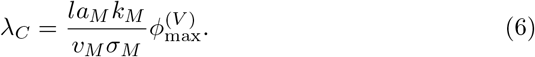

The interpretation of the molecular crowding growth rate limit (6) is straight-forward. The cell content cannot grow faster than the cell capacity to make its membrane.

## 4. Maximum growth rate

Now we stitch everything together taking into account that the protein machinery ℰ_*M*_ is synthesized by the ribosomes. When both ribosomal and ℰ_ℳ_ RNAs are present, the growth rate of the ribosomal and ℰ_ℳ_ proteome contents is given by

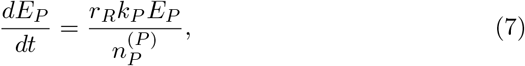

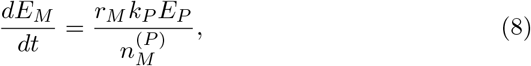

where *r*_*R*_ and *r*_*M*_ (*r*_*R*_ + *r*_*M*_ = 1) are the ℰ_*P*_ and ℰ_*M*_ mRNA fractions. In turn, the cell surface dynamics is described by equation (3). The solution of this system of equations is the exponential growth

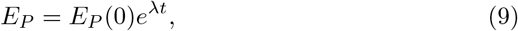

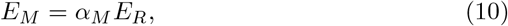

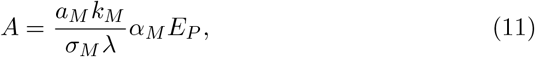

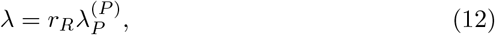

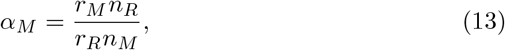

If we stop here, then the obvious strategy to maximize the growth rate *λ* is to maximize the ribosomal RNA fraction *r*_*R*_. For an optimal use of the proteome the cell should allocate as much as possible proteome to the ribosomes. However, that is not the full story. The cell must synthesize its membrane as well and the minimum amount of membrane is constrained by molecular crowding. The exponential growth must satisfy the volume fraction constraint

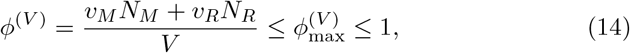

Assuming *V* = *lA* and *v*_*M*_ */n*_*M*_ ≈ *v*_*R*_*/n*_*R*_, equation (14) implies the upper bounds

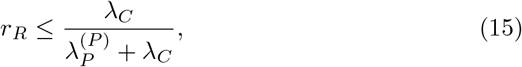

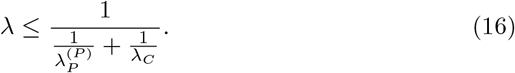

The maximum growth rate is equal to the geometric mean between the protein self-replication rate 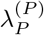 and the maximum surface growth rate *λ*_*C*_ consistent with molecular crowding. The smallest of *λ*_*i*_ will tend to dominate the maximum growth rate. This result was first reported in [16], albeit with an imprecise calculation of *λ*_*C*_. In Ref. [16] I associated *λ*_*C*_ with a non-metabolic protein component whose content scales with cell volume. Here, I am clarifying that the correct protein component is made of the protein complexes responsible for the synthesis of the cell membrane.

## 5. Flux balance analysis

To fully understand metabolism and made accurate quantitative estimates we must resort into full scale models accounting for all the metabolic demands of cells. The simplest approach is to specify the cell composition from experimental measurements and deduce the metabolism that would be consistent with that composition at different growth rates. That is the foundation of flux balance models [17, 18]. The measured cell protein composition can be used to constraint the protein allocated to metabolic enzymes, including ribosomal protein. This is by now a standard practice in the field [19, 20]. In turn, the measured cell composition reflects the relative abundance of cell membrane and enclosed cell components as measured.

What about the self-replication and self-containment constraints described above? Are they taken into account? To answer these questions let’s write down the basic flux balance equations. Let *W* be the cell dry weight, *b*_*i*_ the moles of biomass component *i* per unit of cell dry weight and *E*_*i*_ the total amount of enzyme ℰ_*i*_ responsible for the biosynthesis of biomass component *i*. For example, *i* = *P* is the proteome component and *i* = *M* the cell membrane component. In steady state growth the rate of synthesis balances the growth demand

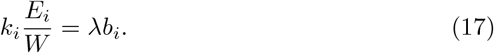

From this equation we can deduce the proteomic fraction 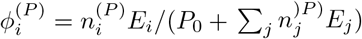 contained in enzyme ℰ_*i*_, where *P*_0_ is the protein content not associated with enzymes and 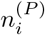 is the number of protein precursors (amino acids) per unit of ℰ_*i*_. From the equation above it follows that

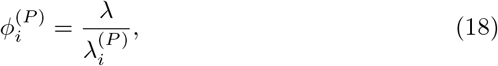

where

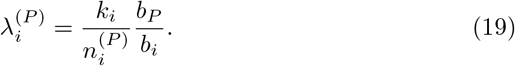

Finally, from the obvious proteomic mass fractions constraint Σ_*i*_ *ϕ*_*i*_ = 1 we obtain

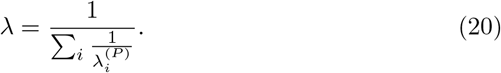

The growth rate is equal to the geometric mean of the growth rates *λ*_*i*_ associated with the biosynthesis of each biomass component *b*_*i*_. This result is equivalent to Theorem 7 in [21].

In particular, for the proteome biomass component (19) simplifies to

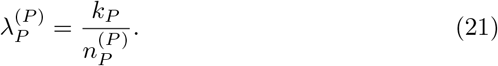

That is the protein self-replication growth rate discussed above (2).

### 5.1. Uncovering the confinement constraint

To uncover the confinement constraint we need to work a bit more. The limited growth rate associated with cell membrane biomass component is

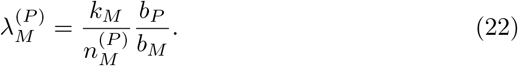

Here *b*_*M*_ is the amount of the membrane precursor per unit of cell dry weight. *b*_*P*_ and *b*_*M*_ can be calculated from the equations

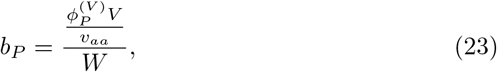

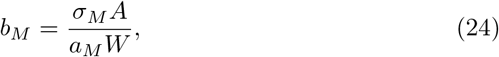

where 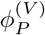 is the cell volume fraction occupied by proteins and *v*_*aa*_ the amino acids molar volume. Substituting (23) and (24) into (22) and taking into account that 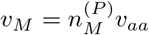 and *A* = *V/l* we obtain

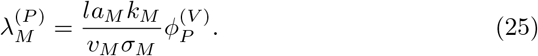

Comparing this result with the equation for *λ*_*C*_ (6) and taking into account that 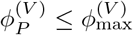 we arrive to the upper bound

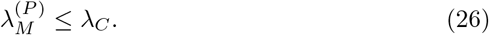

The growth rate limit associated with the synthesis of cell membrane precursors is bound by the maximum growth rate allowed by the confinement constraint.

Finally, substituting the growth rates (21) and (27) on the geometric mean equation (20) we arrive to the upper bound

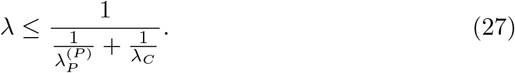

This result recapitulates our earlier calculation leading to equation (16). Here we have neglected the contribution of other components such as RNA, DNA and lipids. Adding those components will made the upper bound tighter.

### 5.2. What about RNA fractions?

RNA is another mayor component of the cell dry weight. It is worth asking what can be concluded if we focus on the RNA content of enzymes. That is the case of ribosomes, with a composition of about half protein and half ribosomal RNA (rRNA). The RNA fraction included in enzyme ℰ_*i*_ is

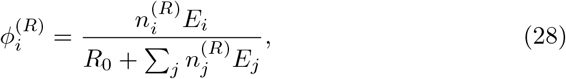

where *R*_0_ is total RNA not contained in enzymes and 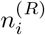 the number of RNA precursors (ribonucleotides) per enzyme ℰ_*i*_. Using equation (17) we obtain

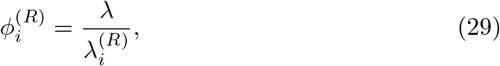

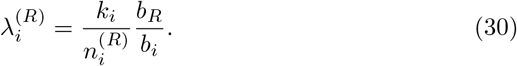

Finally, from the obvious RNA fraction constraint 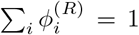 we arrive to another expression for the growth rate

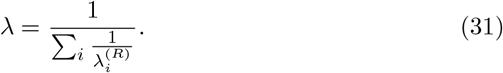

Ribosomes, the protein synthesis machinery (*i* = *P*), are the most abundant RNA containing enzyme and therefore (31) can be approximated by

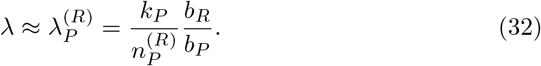

However, since we cannot put a bound to the right hand side, this equation just tells us that *k*_*P*_ *b*_*R*_*/b*_*P*_ grows linearly with the growth rate. Working with the RNA fractions does not lead to further inside regarding the maximum growth rate.

## 6. Discussion

I hope the material presented above clarifies the interplay between self-replication and self-confinement in determining the maximum growth rate of a cell. Self-replication has been extensively studied and it was analyzed here for the sake of completeness. The main conclusion is: (i) Because the replication machinery produces itself, then the production of anything else reduces the growth rate.

The impact of self-confinement is less understood however. Here I have shown that: (ii) Self-confinement implies the geometrical constraint that the cell volume cannot grow faster than the volume included by the cell cell surface. In turn it implies that the macromolecular components making up the cell surface need to be synthesized fast enough to cope with the growth of the volume occupied by all macromolecules within the cell.

The synthesis of the macromolecular machinery responsible for the synthesis of the cell surface will deviate resources from self-replication, those reducing the growth rate. To achieve the maximum growth rate the cell needs to be contained within the minimal volume and shape that would require the minimal surface area. That minimal volume is constrained however by molecular crowding. As a corollary, to achieve the maximum growth rate the macromolecules in a cell should be as crowded as possible. Selection for maximum growth rate selects for molecular crowding.

### 6.1. Modeling cell growth

When modeling cell growth, we should distinguish the scenarios where we are or are not fixing the biomass vector. Models with a fixed biomass vector are satisfying the self-confinement constraint, at least in a first approximation. In reality the cell always satisfies the self-confinement constraint and the biomass composition was derived from cell measurements. That means that the ratio between the surface and the intracellular macromolecular components is balanced, in a first approximation.

It is an approximation because the model may allow for a variable fraction of proteins that localize to the cell membrane and the intracellular volume, albeit with a fixed total protein abundance. The model composition may deviate from the one where the biomass measurements were taken from. In such case the self-confinement constraint may be violated.

In contrast, the self-confinement constraint is essential when developing first principles models that aim to predict the cell biomass composition itself. In this context, we should bear in mind that molecular crowding affects the kinetic constants of biochemical reactions. A full metabolic model is more complicated than what presented here (see for example Ref. [22]).

### 6.2. Modeling the metabolism of non-growing cells

The picture is different in non-growing cells. Here self-replication is by definition irrelevant. What matter most is that non-proliferating cell fulfills some primary function and their resources should be devoted, as much as possible, to that purpose. Hypothesis: In non-proliferating cells the primary complexes responsible for the cell function are at the maximum concentration allowed by molecular crowding. All other support complexes are selected for maximum yield per unit of occupied volume. In non-growing cells self-replication plays no role and self-confinement becomes the key metabolic constraint.

For example in myocytes, the muscle cells, the key component are the myofilaments. They are responsible for transducing the chemical energy from ATP hydrolysis into mechanical work. In turn, the ATP regenerating pathways play a support role. Optimization of myocytes for maximum mechanical work should select for the highest concentration myofilaments and the ATP generating pathway with maximum ATP production per unit of occupied volume. Glucose fermentation to lactic acid has a higher yield of ATP per unit of occupied volume that oxidative phosphorylation [15, 23], providing an explanation for the use of fermentation during intense physical activity.

The full extent of this hypothesis remains to be investigated in other non-proliferating cells.

## Notes

### Competing Interest Statement

The authors have declared no competing interest.

